# Gene-signatures predict biologically relevant dose-response potencies in phenotypic assays

**DOI:** 10.1101/799106

**Authors:** Steffen Renner, Christian Bergsdorf, Rochdi Bouhelal, Magdalena Koziczak-Holbro, Andrea Marco Amati, Valerie Techer-Etienne, Ludivine Flotte, Nicole Reymann, Karen Kapur, Sebastian Hoersch, Edward J. Oakeley, Ansgar Schuffenhauer, Hanspeter Gubler, Eugen Lounkine, Pierre Farmer

## Abstract

Multiplexed gene-signature-based phenotypic assays are increasingly used for the identification and profiling of small molecule-tool compounds and drugs. Here we introduce a method (provided as R-package) for the quantification of the dose-response potency of a gene-signature as EC_50_ and IC_50_ values.

Two signaling pathways were used as models to validate our methods: beta-adrenergic agonistic activity on cAMP generation (dedicated dataset generated for this study) and EGFR inhibitory effect on cancer cell viability. In both cases, potencies derived from multi-gene expression data were highly correlated with orthogonal potencies derived from cAMP and cell growth readouts, and superior to potencies derived from single individual genes.

Our results show that gene-signature potencies are a novel valid alternative to conventional readouts for compound potency quantification, in particular in scenarios where no other established readouts are available.

## Introductions

Gene expression signatures are widely used in the field of translational medicine to define disease sub-types [1], severity [2] and predict treatment outcome [3]. Bridging this technology to early drug discovery was previously proposed years ago [4, 5] but its prohibitive costs limited this approach. The recent advancement of massively parallel gene expression technologies such as RASL-seq [6], DRUG-seq [7], QIAseq [8, 9], PLATE-seq [10], or LINCS L1000 [11] are now transforming the field of compound profiling, enabling larger scale profiling and screening experiments at a more affordable cost [12–17].

In drug discovery, dose-response experiments enable researchers to compare the efficacy of various compounds to modulate biological processes of interest, finding doses for animal and human experiments and estimating windows to off-target and toxic effects. Multiple statistical methods are reported for the identification of individual genes with a dose dependent effect from dose-response gene expression data [18–23]. However, in the case of multivariate gene expression profiling there are no generally accepted methods to estimate the key pharmacological efficacy variables EC_50_ (compound concentration of half-maximal activating effect) and IC_50_ (compound concentration of half-maximal inhibitory effect) from multi-parametric readouts.

Connectivity Map (CMap) established the concept that compounds with similar mode of actions (MOAs) are highly similar in their differential expression profiles over many genes [4, 11, 24]. We postulate that this concept can be applied for quantifying compound potencies based on compound/pathway specific gene expression signatures. This work aims at defining and comparing several multivariate statistical summaries to enable classical compound potency estimation. In this study, we focus mainly on methods measuring the similarity of gene-signature changes relative to a gene-signature induced by an active control compound, representing a defined phenotype of interest, e.g. a tool compound for a target or pathway of interest. The overall principal relies on assessing the similarity of a compound-induced gene-signature profile relative to the one generated by an active control compound; hence, the AC profile will anchor all other measurements in the form of a global reference.

The different similarity methods explored in this paper differ by their approach to assess either the direction of the effect (as example by the geometric angle (cosine) to the AC; referred as direction-based methods) and / or by how the magnitude of the effect is assessed (e.g. Euclidean distance to the NC, referred as magnitude-based methods). Combined, the two measures quantify the strength and direction of a phenotypic effect (see Fig. 1 and Table 1). Methods referred to as direction&magnitude-based combine both types of information into a single measure.

**Table 1:**
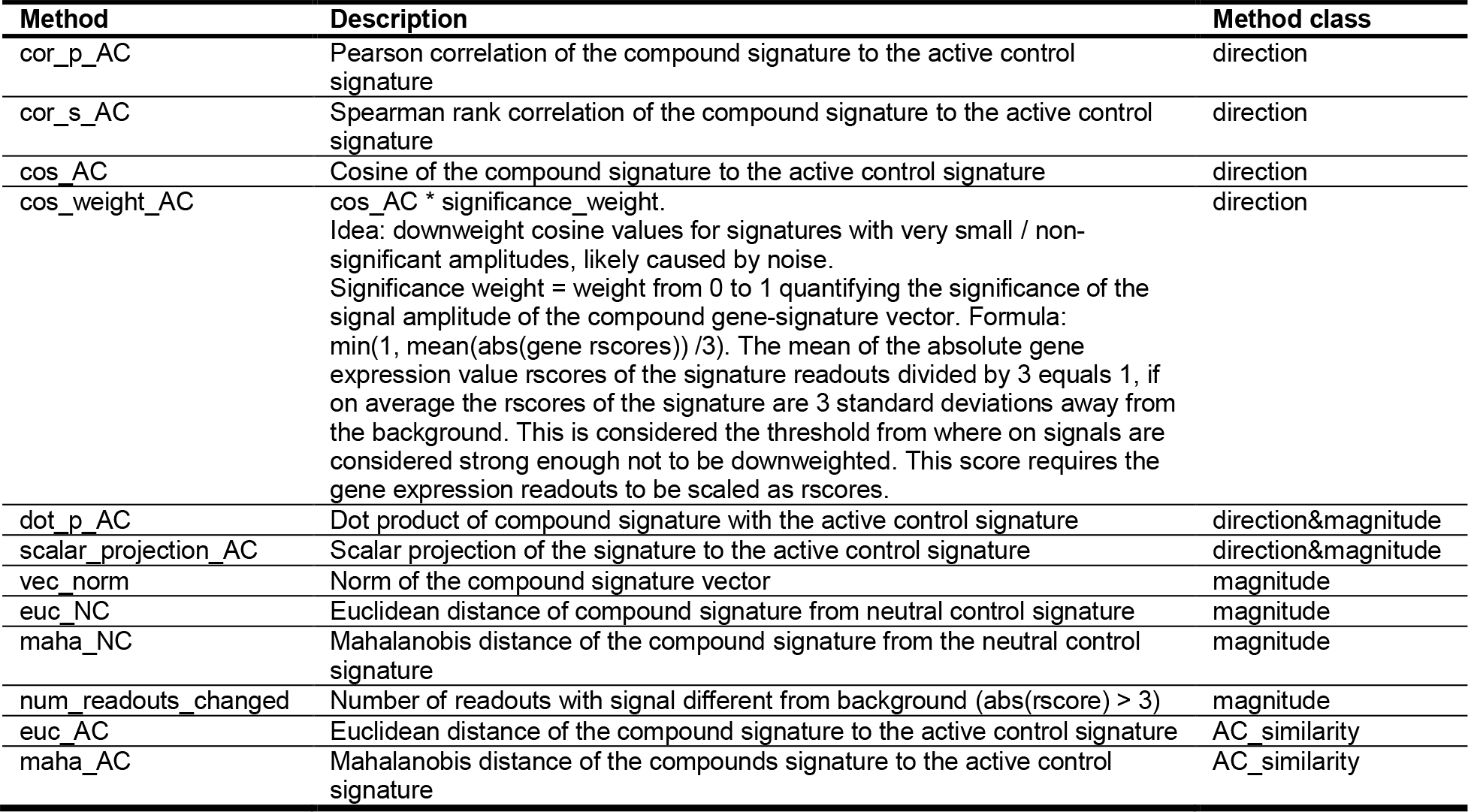
Overview over gene-signature quantification methods.

**Figure 1.**
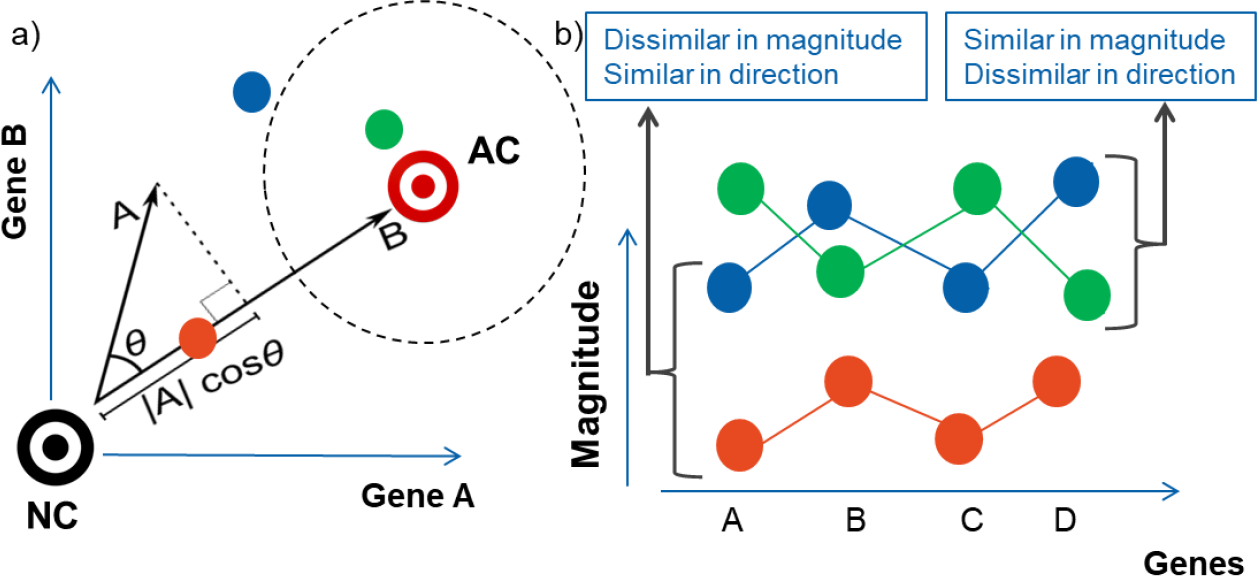
Introduction to gene-signature quantification methods. a) Within the manuscript, we consider methods measuring the similarity of gene-signature changes relative to an active control (AC) or a neutral control (NC). b) Two main characteristics of signature similarity can be distinguished: similar changes in magnitude or similar changes in the direction of the gene expression. The magnitude can be interpreted biologically as the efficacy, while the direction emphasizes the direction of the change of the phenotype, e.g. different pathways might result in different directions of changes in gene expression.

For this study, two well-characterized biological pathways with multiple well-characterized ligands were selected: the beta-adrenergic receptor pathway for which we generated experimental biological data for this manuscript, and the EGFR pathway, which is publicly available through the LINCS L1000 project [11]. For the beta-adrenergic pathway we used cAMP EC_50_s as functional orthogonal readout [25]. For practical purposes, we had to measure a small set of biologically relevant genes, instead of the full transcriptome like in CMap. RNA-seq was used to determine a beta-adrenergic receptor specific gene-signature that was subsequently used to quantify compound potencies on the level of gene expression. The L1000 assay is a panel of ca. 1000 measured genes, which are used to infer the differential gene expression of a total of ca. 13k genes. This allowed us to benchmark our methods using all L1000 genes, and subsets thereof specific for EGFR signaling or cell proliferation. The IC_50_s calculated from gene expression were compared to compound potencies measuring the inhibition of cell growth rate (GR_50_) [26].

Our results demonstrate that gene-signature-based compound EC_50_ and IC_50_ values estimated with multi-variate gene-signatures are highly related to potencies inferred with relevant but independent reference readouts. Therefore, we expect that these methods will find a wide application in gene-signature based assays in the near future. All methods in Table 1 and an EC_50_ and IC_50_ fitting method are made available in the R-package mvAC50 on github [https://github.com/Novartis/mvAC50].

## Results

### Generation of the beta-adrenergic receptor dataset

Vitamin-D3 differentiated THP1 cells were chosen as an experimental model for its sensitivity to beta agonists over a large dynamic range of compound concentrations and the ease of measuring cAMP [27]. To identify a gene-signature for beta agonists, a series of RNA-seq experiments were performed on THP1 cells sampled at baseline and after four hours stimulation with adrenaline, noradrenaline or isoproterenol.

Genes differentially expressed over all three treatments were identified, and prioritized for large fold change and high expression levels, for independent qPCR validation (Supplementary Fig. 1a). Our internal compound screening setup allows us to simultaneously multiplex the measurement of eight genes. Two independent sets of seven genes were defined from 14 qPCR validated genes (Supplementary Table 1, Supplementary Fig. 1b) with the eighth gene per set (TBP) serving as a baseline house keeper gene. For our analysis, we considered the two sets of genes as two independant signatures. Not all of these 14 identified genes produced a detectable signal in the QuantiGene Plex technology due to decreased sensitivity of this method compared to qPCR (Supplementary Fig. 1b). The two sets of genes contain respectively three (CD55, DOCK4, and NR4A1) and five genes (PDE4B, SGK1, THBS1, TOB1 and VEGFA) responding consistently to 10uM of isoproterenol.

### Comparison of EC_50_s from single genes, gene-signatures, and cAMP

A total of 21 beta agonists (Supplementary Table 2) covering a wide range of potencies (<10pM to ca. 5uM), were chosen for this study. Other cAMP modulators were also included in this compound set: the histamine receptor H3 antagonist N-alpha-methylhistamine and the adenylyl cyclase activator forskolin. As additional control, we added the beta-1 antagonist CGP-20712A, which, as expected, failed to increase cAMP levels. All compounds were measured in dose-response mode in the cAMP assay and for both gene signatures. An overview of dose-response curves of the genes is shown in Supplementary Fig. 2. The gene-expression data, derived gene-signature scores, and fitted EC_50_s are presented in Supplementary Tables 3 and 4.

The relationship of EC_50_ values derived from genes and gene-signatures compared to cAMP-derived EC_50_s depends on the gene-signature methods used. Representative examples for method classes are shown in Fig. 2a. (all methods and genes are shown in Supplementary Fig. 3). The EC_50_s derived from direction-based methods cor_p_AC and cos_weight_AC are found almost entirely within a window of one log unit around the cAMP-derived EC_50_s, which is very close considering the different incubation times and the different locations of the readouts in the adrenergic signaling pathway (gene expression vs cAMP). In contrast, the EC_50_s derived from gene-signature methods containing magnitude information (scalar_projection_AC and vec_norm) and EC_50_s from the individual genes NR4A1 and THBS1 are almost all more than one log unit above the cAMP-derived EC_50_s. The ranking of cAMP potencies is not preserved as well (e.g. Spearman correlation for scalar_projection_AC to cAMP = 0.32).

**Figure 2:**
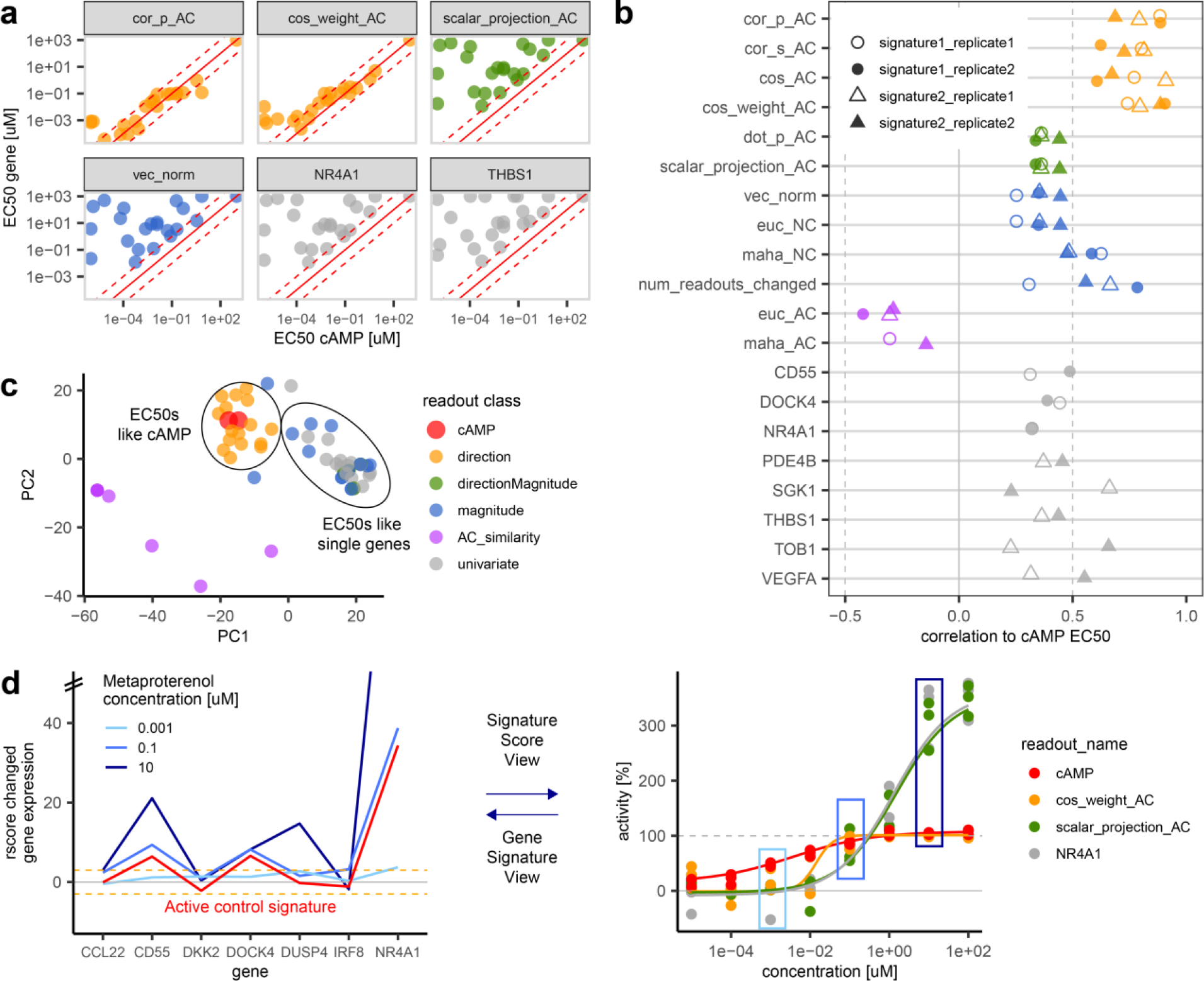
Comparison of EC_50_s from gene-signatures, single-genes and cAMP for the beta agonists dataset. a) Example of gene and gene-signature EC_50_s from representative methods compared to cAMP EC_50_s. The dashed red lines indicates one log unit above and below the red line of equality. The shown gene and gene-signature EC_50_s are from signature one, except THBS1 from signature two. The shown data is from replicate two. Axes are log10 transformed. b) Correlation of gene-signature and single-gene EC_50_s with cAMP EC_50_s. c) PCA of the cAMP, gene, and gene-signature summary methods logged EC_50_s of all compounds in the dataset. Colors of a), b), and c) are according to the definition in c). d). Dose dependent change of the genes in the gene-signature (left panel, with y-axis values > 50 not shown, orange dashed lines at three rscores indicating significant changes from the background), compared with the dose dependent change in gene-signature summary score methods and cAMP for metaproterenol (right panel, boxes colored according concentrations shown in left panel, dashed grey line at 100% activity).

A performance overview of all genes and gene-signature methods is given in Fig. 2b. The similarity between gene or gene-signature derived EC_50_s with cAMP derived EC_50_s over all tested compounds is quantified by the Pearson correlation between logged EC_50_s. Most methods within one method-class performed equally well. While direction&magnitude and magnitude-based methods showed no significant difference to individual genes (TukeyHSD test with p-val < 0.05), direction-based methods performed significantly better than the other methods with Pearson correlations ranging between 0.6 and 0.9. All other method classes showed mean Pearson correlations < 0.5. The AC_similarity method performed significantly worse relative to others (only negative correlations).

The relationship between all gene-signature methods, single genes and cAMP EC_50_s is shown by a principle component analysis (PCA) projection of the dataset (Fig. 2c). Each data point represents the vector of logged EC_50_s calculated by one method (for one replicate and one gene-signature) of all compounds in the dataset, Methods generating similar EC_50_s are projected close to each other. The PCA projection confirms that direction methods cluster together with the cAMP EC_50_s, and all EC_50_s containing magnitude information cluster together with single gene EC_50_s. As mentioned above, the AC_similarity methods are outliers relative to the two major clusters.

Fig. 2d visualizes the expression levels of the individual genes over compound concentrations (left panel) and the resulting dose-response curves of derived multivariate EC_50_ methods (right panel). Increasing concentrations of metaproterenol result in increasing expression of the genes of the gene-signature. While the shape of the gene-signature remains similar to the active control signature (isoproterenol [10uM], red line), the magnitude of the metaproterenol signature exceeds the AC signature with increasing concentrations (left panel). The observed difference in gene expression magnitude between high concentrations of metaproterenol and the active control signature is only captured by metrics that make use of this information (Fig. 2d, right panel, green line). It is important to note that the difference between methods does not only lead to different maximal effect plateaus of the dose-response curve, but also to different EC_50_ values of the fitted curves.

The increase in gene expression beyond the active control also explains why AC_similarity methods cannot work in this scenario: the maximum similarity between compounds and AC signature is reached at identical magnitudes of both signatures. Both lower and larger magnitudes result in less similar signatures, resulting in bell shaped curves.

### EGFR inhibitors dataset from L1000

For the L1000 EGFR (“Epidermal growth factor receptor”) inhibitor dataset, we selected a set of eight EGFR inhibitors measured in six-point dose-response in MCF10A cells after 3h and 24h incubation time [11]. As reference univariate readout, the corresponding growth rate inhibition GR_50_ measured after three days was used [26]. As the LINC technology reported 12,717 genes, it was possible to test several gene-signatures: (1) a published EGFR signature [28], and (2) a published cell proliferation gene-signature [3], further referred to by the gene name “Targeting protein for Xklp2” (TPX2). As a third biologically unbiased gene-set, all genes from L1000 were used for comparison. We also investigated the performance of single gene measurements, for which we chose the 20 genes from each of the three signatures with the strongest response to the active control (gefitinib at 3.33uM).

Like with the beta agonist pathway data, gene-signature IC_50_s of the EGFR inhibitors corresponded well to the reference GR_50_s (Fig. 3a for representative readouts, all results in Supplementary Fig. 4-6). Results show a strong influence of the incubation time. At 24h all shown gene-signature methods over all three gene-signatures resulted in IC_50_ vs GR_50_ correlations >= 0.88, except scalar_projection_AC and vec_norm with the TPX2 gene-signature resulting in slightly lower correlations each of 0.68. The individual single gene IC_50_s at 24h incubation showed more variance, with Pearson correlations ranging from −0.36 with the TPX2 signature to 0.9 with the EGFR signature. The individual genes from the EGFR signature resulted in the highest median correlation of 0.88. Two very similar median correlations of 0.68 and 0.69 were found for the individual genes of the TPX2 signature and from all L1000 genes, confirming the lower biological relevance for the EGFR pathway of the latter signatures. Even though all three gene-signatures contained individual genes that correlated very well with the GR_50_s (> 0.9), all of them also contained genes with correlations to GR_50_s < 0.5, few even around 0. It is not clear how one could reliably distinguish more relevant from less relevant genes in the absence of another orthogonal reference-readout like the GR_50_s.

**Figure 3.**
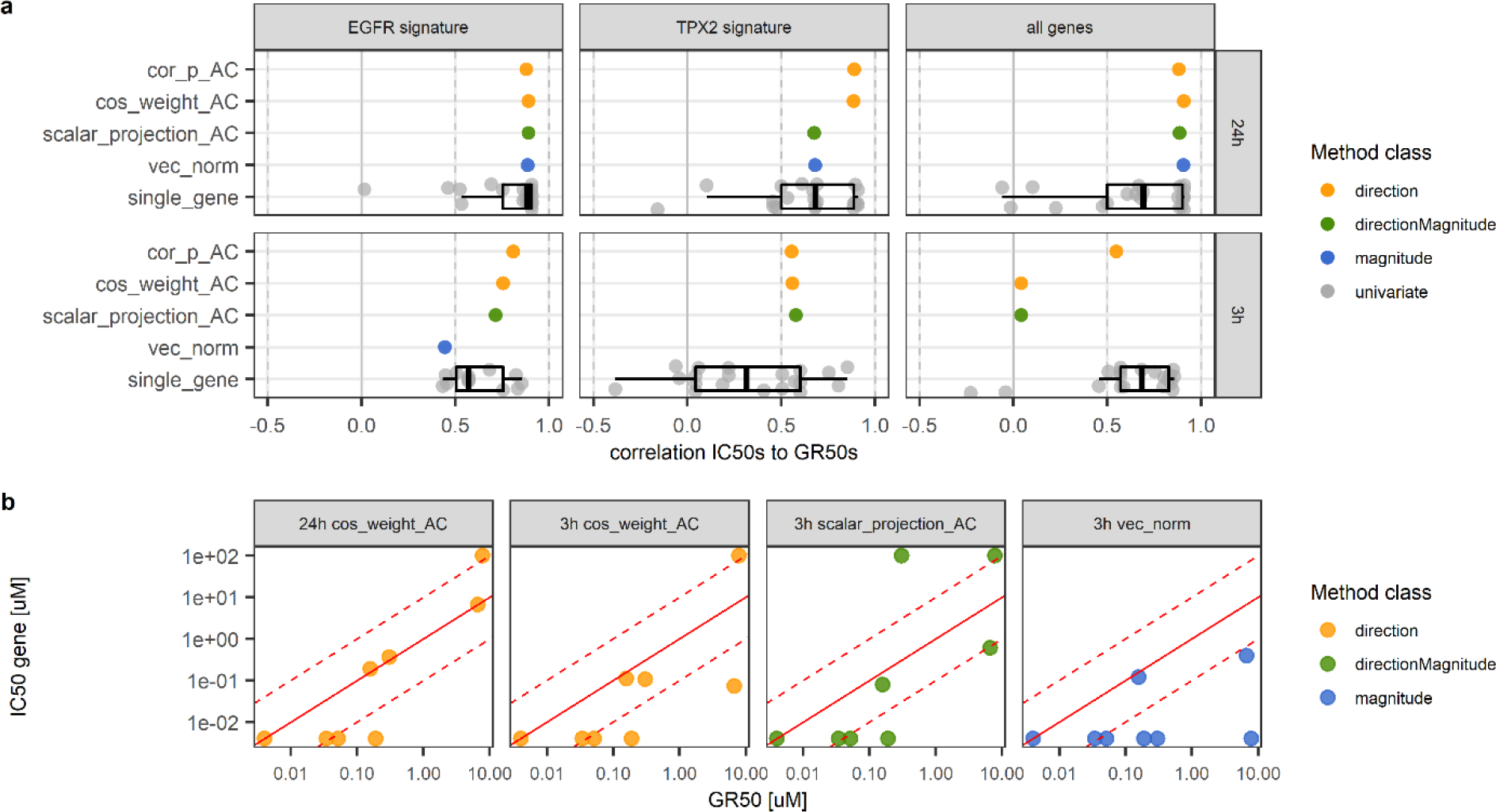
Comparison of gene and gene-signature IC_50_s to growth rate inhibition GR_50_s of EGFR inhibitors. a) Pearson correlation of representative methods and 20 individual gene IC_50_s to GR_50_s. b) Comparison of representative EGFR gene-signature IC_50_s in MCF10A vs. GR_50_s. The dashed red lines indicate one log unit above and below the red line of equality.

At 3h incubation time, differences between methods and gene-signatures are more pronounced, showing highest correlations for direction-based methods with the EGFR signature (both above 0.75). Again individual genes show a wide distribution of results ranging from −0.38 for TPX2 to ca. 0.85 for all three gene-sets. Like with the beta agonists, the values of gene-signature IC_50_s are very close to the values from GR_50_s and more than 50% of the gene-signature IC_50_ values are within a one-log-unit window to the GR_50_s (Fig. 3b).

## Discussion

The two main contributions of this work are: (1) the development and validation of an analytical framework for calculating compound potency based on multivariate readouts and (2) the provision of an open-source R-package to facilitate the application of our methods on new data by the scientific community.

The principal of this framework is to first summarize the information contained in multiple-genes into a single value and then pass it into a logistic function for potency estimation. The optimal metrics were selected based on their degree of concordance with compound potencies estimated with standard readouts (cAMP/GR_50_).

The fact that IC_50_/EC_50_ potency measurements are specific to a given biologic process (cAMP, gene expression, cell viability), and not a general property of the compound, is a potential challenge for comparing methods. However, choosing experimental models where gene expression is closely linked to pathway activation provides us confidence in our working model. The conservation of the compounds potency rank-order regardless of using gene expression or standard readouts supports our premise. Indeed, very close potency relationship (Pearson correlations up to 0.9) were observed for reference potency values (cAMP, GR_50_) upstream (cAMP) and downstream (GR_50_) of the gene expression readout, and independent of very different compound incubation times of readouts. The assessments of optimal methods was not influenced by gene-signature composition. Indeed, all signatures used in this work were previously reported, or constructed independently of the screening datasets.

Of the five methodological classes of metrics: (1) direction-based, (2) distance based (magnitude) to the NC, (3) distance based (magnitude) to the AC, (4) magnitude and direction-based and (5) single genes, results show that magnitude-based methods to the AC clearly underperformed to other methods while direction-based methods performed consistently well in the two explored datasets. We did not find large differences in the performance of the methods within a single method class in these two datasets. Yet we recommend cos_weight_AC for direction-based methods due to its ability to down-weight signal with very small magnitude. To our surprise, adding information about the magnitude of the gene expression did not improve the results.

To this date, there is still very limited data available in the public domain that enables the comparison of multivariate EC_50_/IC_50_ with standard readouts, hence it is impossible to generalized current findings to future situations. Nonetheless, with the raise of novel sequencing methods that enable low to medium throughput compound screening based on hundreds to thousands of genes, the need for multivariate potency estimation will be strong.

Finally, our work enables the of use gene-signatures as screening readouts and biomarkers throughout all stages of research from early cell line experiments, to animal models and clinical studies. Using the same readout will in many cases contribute to increased biological relevancy at all stage of the drug discovery process. Similar multiplexed readouts like the data from cell painting or metabolomics [29, 30] might also benefit from our multiplexed potency methods.

The publication of the first dedicated dataset to investigate the quantification of the dose-response based on gene-signatures together with the first analysis of such data and the publication of an R-package providing the methods for such analyses will enable the further exploration and application of these methods by the scientific community. The algorithms and datasets used in the publications are available in the R-package mvAC50 from https://github.com/Novartis/mvAC50.

## Abbreviations

CMap: Connectivity Map
EC_50_: Compound concentration of half-maximal activating effect
IC_50_: Compound concentration of half-maximal inhibitory effect
AC_50_: Compound concentration of half-maximal effect, independent of curve direction. General term including both EC_50_ and IC_50_
GR_50_: Compound concentration of 50% reduction of the GR values, where the GR value is the ratio between growth rates under treated and untreated conditions normalized to a single cell division
AC: active control
NC: neutral control

## Methods

### THP1 cells

Human promonocytic THP-1 cells (TIB-202, ATCC) were cultured at 37°C/CO_2_ in medium (Hepes (72400-054, Life Technologies), with 10% FBS (2-01F16-I, Amimed/Bioconcept), 1% Pen/Strep (15140-122, Life Technologies), 1mM Sodium Pyruvate (11360-039, Life Technologies), 2mM L-Glutamine (25030-024, Life Technologies), 0.0mM Mercaptoethanol (31350-010, Life Technologies)). Before compound treatment and for all experiments, the THP1 cells were differentiated with 100nM Vitamin D_3_ (Biotrend Chemicals AG, Switzerland, Cat. No. BG0684) for 3 days at 37°C/CO_2_.

### cAMP HTRF assay

The assay was run using the Cisbio cAMP dynamic 2 Kit (62AM4PEB), in white 384well-plates BioCoat #354661, with 20,000 cells/well in 10μL/well HBSS/HEPES/IBMX. Isoproterenol [10uM] was used as active control. Cells were incubated with compounds for 20 min. at 37°C in HBSS/HEPES, in the presence of the Phosphodiesterase (PDE) inhibitor IBMX. Then, cells were lysed and the amount of generated cAMP was quantified by HTRF (Homogeneous Time Resolved Fluorescence).

### Beta agonists gene-signature

RNASeq experiments were done comparing untreated cells with a treatment with isoproterenol, adrenaline or noradrenaline for 4h in THP1 cells.

qPCR was run in THP1 cells for 4h incubation time with isoproterenol and formoterol at 1, 10 and 100 nM. Total RNAs were isolated with MagMAX™-96 Total RNA Isolation Kit (Ambion ref#AM1830), and cDNA was made using a cDNA Synthesis Kit (Applied Biosystems™ Ref#4368813) RT-PCRs were performed in 384-well plates on an AB7900HT cycler (Applied Biosystems) using specific TaqMan probes (Applied Biosystems). Housekeeper normalization was done relative to the one of the three genes GAPDH, PPIB or TBP, which had the most similar expression level to the gene of interest, according to our DMSO qPCR data. All measurements were done in quadruplicates.

### QuantiGene Plex assay

Gene expression changes were measured using a customized QuantiGene Plex assay (Thermo Fisher Scientific).

Two different eight-gene-signatures were designed (obtained from Thermo Fisher Scientific), as the internal QuantiGene process was set up to handle custom-designed signatures of eight genes. Each of the eight-gene-signatures consisted of seven target genes responding to cAMP and one housekeeper gene (TBP).

Measurements were done in THP1 cells. Compounds were measured in six replicates on the same day on different plates, and the procedure was repeated on another day using three replicates on different plates (referred to as biological replicates in the manuscript).

For the assay, 100,000 cells were seeded in a volume of 20uL in each well of a 384 well plate (Greiner PP V bottom 781280). Compounds were added in serial dilutions of 1:10 (200nL volume added per well) with maximal compound concentrations of 100uM. After 4h incubation, cells were lysed with QuantiGene lysis mixture (10uL), and after 2 min, stored at −80°C.

Targeted mRNA transcripts were captured to their respective beads by combining lysis mixture (5uL), blocking reagent (2uL), probe mix (1.125uL), water (11.25uL), and magnetic beads (0.3 uL; 500 beads/region/uL) and incubated overnight.

Signal amplification via branched DNA is added by sequential hybridization of 2.0 Pre Amplifier biotinylated label probe, and binding with Steptavidin-conjugated Phycoerythrin (SAPE). For this purpose, each 15uL/well pre-amplifier, amplifier and label probe & SAPE were added after washing followed by 1h incubation at 50°C and multitron shaking 300rpm 1h.

The amount of RNA in 90uL of probe was quantified using a Luminex Flexmap 3D instrument (Luminex). The identity of the mRNA is encoded by the hybridized Luminex beads, and the level of SAPE fluorescence is proportional to the amount of mRNA transcripts captured by the respective beads.

### QuantiGene Plex data processing

The raw readout of the assay was processed as follows:

1. Fold change = 50 * log_2_ (mRNA count / median mRNA count for NC well)
2. Rscore = (Fold change for well − median Fold change for NC wells) / MAD (mRNA count for NC wells)
3. HKnorm = Rscore for well − HK_Rscore for well; with HK = housekeeper gene.

### L1000 / GR_50_ dataset

EGFR inhibitors in MCF10A cells were selected as model system, because (1) they showed a strong GR_50_ dynamic range (Dose-response curves visualization http://www.grcalculator.org/grbrowser/.), and

(2) were measured in six concentrations in L1000 (10uM, 3.33uM, 1.11uM, 0.37uM, 0.12uM, 0.04uM).

The L1000 data was obtained in two files (GSE70138_Broad_LINCS_Level4_ZSVCINF_mlr12k_n78980×22268_2015-06-30.gct.gz and GSE70138_Broad_LINCS_Level4_ZSVCINF_mlr12k_n115209×22268_2015-12-31.gct.gz) from NCBI GEO (https://www.ncbi.nlm.nih.gov/geo/query/acc.cgi?acc=GSE70138).

This version of the data contains the changed gene-expression normalized as z-scores relative to the DMSO controls on each plate, a similar normalization procedure to the one performed for the beta-agonists expression data. When multiple probes were measured for the same gene_symbol, the probe with the highest variance was kept, for each timepoint. The gefitinib treatment at 3.33 uM was defined as the active control of the experiment. Compounds, smiles, and inchi_key were downloaded from the LINCS webpage (http://lincs.hms.harvard.edu/db/datasets/20000/).

From the 12,727 genes in the L1000 dataset, two different subsets were selected based on published gene-signatures. An EGFR (entrez gene_id 1956) signature [28] (“EGFR_UP.V1_UP” with 193 genes, “EGFR_UP.V1_DN” with 196 genes) was downloaded from msigdb [31, 32], of which a total of 381 genes could be mapped to the L1000 data. This gene-signature was derived from profiling of MCF-7 cell lines stably overexpressing ligand-activatable EGFR. A TPX2 (entrez gene_id 22974) signature (50 genes, of which 39 could be mapped to L1000) was taken from Farmer et al [3], representing a more general signature for cell proliferation.

The GR_50_ cell viability potency values after three days compound incubation time were also obtained from the LINCS consortium (http://lincs.hms.harvard.edu/db/datasets/20252/results). To make the data more comparable to the fitted IC_50_’s from the gene-signatures, compounds with flat GR_50_ dose-response curves were set to either one log unit above or below the highest or lowest tested concentration, depending whether their fitted GRInf value was larger or smaller than 0.5.

As the files from L1000 and the GR_50_s contained slightly different compound and cell line names, the names were set all to lowercase and whitespaces and “-“ were removed. Eight known EGFR inhibitors afatinib, neratinib, pelitinib, gefitinib, erlotinib, canertinib, lapatinib, and HG-5-88-01 overlapped between the two datasets. The two EGFR/ERBB2 dual inhibitors neratinib and afatinib were considered as EGFR inhibitors for this study (even though they are annotated as ERBB2 inhibitors in the LINCS nominal target annotation).

### Dose-response (DRC) fitting

Four-point parametric logistic fits were calculated with an R function included in the mvAC50 R-package [https://github.com/Novartis/mvAC50]. The fitting algorithm was adopted from our in-house HTS analysis software Helios [33]. The fits were constrained to A0 and Ainf (minimal and maximal fitted activities) between −50% and 500% of the active control effect, respectively, and a hill slope between 0.1 and 10. The IC_50_s or EC_50_s were constrained to one log unit above and below the experimentally measured range of concentrations, (for the beta agonists ranging from 0.00001uM to 100uM, and for the L1000 data ranging from 0.04uM to 10uM)

In the case of constant fits, IC_50_ or EC_50_ values one-log unit above or below the range of tested concentrations were assigned to the compounds to be able to use those data points as well in the correlation of calculated potencies to the reference potencies. Depending on whether the Amax of the constant fit was below or above 50%, a potency of either one log unit below or above the tested concentration range was assigned. Fitted AC_50_s with Ainf values < 50% were set to one log unit above the highest tested concentration as well, assuming that the observed effect is not caused by the same mode of action as in the active control.

In parallel to the four-point parametric fit and constant fits, a nonparametric fit was also calculated and compared to the other fits, to allow for more unusual curve shapes, e.g. bell shaped curves. For these fits the reported potency is the concentration at which the fit crosses the line of 50% activity. The decision for the reported fit and potency was done as follows: If the non-parametric fit resulted in r2 < 0.5, the data was considered as not suitable for curve fitting and assigned as constant fit. If the curve had a bell-shape, the nonparametric potency was reported. If parametric fits had r2 < 0.5 or the absolute (amin-amax) < 30, a constant fit was reported as well, where amin and amax correspond to A0 and Ainf within the measured concentration range. For the remaining curves (the majority) parametric potencies were reported.

cAMP EC_50_s were fitted with the same algorithm and settings, to ensure a higher consistency in the data. The fitted cAMP EC_50_s were in agreement with the fits generated by the biologists who ran the assays. For the GR_50_ dataset this approach was not feasible, as no raw data was available, and the GR_50_ algorithm was claimed to be superior to four-point parametric fits of the same data [26].

## Acknowledgment

We would like to acknowledge Stan Lazic, Xian Zhang, and Jeremy Jenkins for helpful discussions about the concept of multivariate AC_50_s, Wendy Broom, Elaine Donohue and Jacques Hamon for help with the QuantiGene assay, Magalie Mathies for help with setting up the THP-1 assays, Pierre Rigo, Thomas Hoerter, Cornelia Mouzo and Valerie Heidinger for production of THP-1 cells, Ioannis Moutsatsos for help with the QuantigGene analysis pipeline, and Pascale Anderle for referring Andrea Amati as NIBR intern for this project.

## Supplementary Information

**Supplementary Table 1:** Genes selected for quantification of beta agonist potencies.

| Gene symbol | Gene id | Name | Set | Comment |
| --- | --- | --- | --- | --- |
| CCL22 | 6367 | C-C motif chemokine ligand 22 | 1 |  |
| CD55 | 1604 | CD55 molecule (Cromer blood group) | 1 |  |
| DKK2 | 27123 | dickkopf WNT signaling pathway inhibitor 2 | 1 |  |
| DOCK4 | 9732 | dedicator of cytokinesis 4 | 1 |  |
| DUSP4 | 1846 | dual specificity phosphatase 4 | 1 |  |
| IRF8 | 3394 | interferon regulatory factor 8 | 1 |  |
| NR4A1 | 3164 | nuclear receptor subfamily 4 group A member 1 | 1 |  |
| TBP | 6908 | TATA-box binding protein | 1 | Housekeeper |
| NR4A3 | 8013 | nuclear receptor subfamily 4 group A member 3 | 2 |  |
| PDE4B | 5142 | phosphodiesterase 4B | 2 |  |
| PPARGC1B | 133522 | PPARG coactivator 1 beta | 2 |  |
| SGK1 | 6446 | serum/glucocorticoid regulated kinase 1 | 2 |  |
| THBS1 | 6908 | TATA-box binding protein | 2 |  |
| TOB1 | 7057 | thrombospondin 1 | 2 |  |
| VEGFA | 10140 | transducer of ERBB2, 1 | 2 |  |
| TBP | 7422 | vascular endothelial growth factor A | 2 | Housekeeper |

**Supplementary Table 2:**
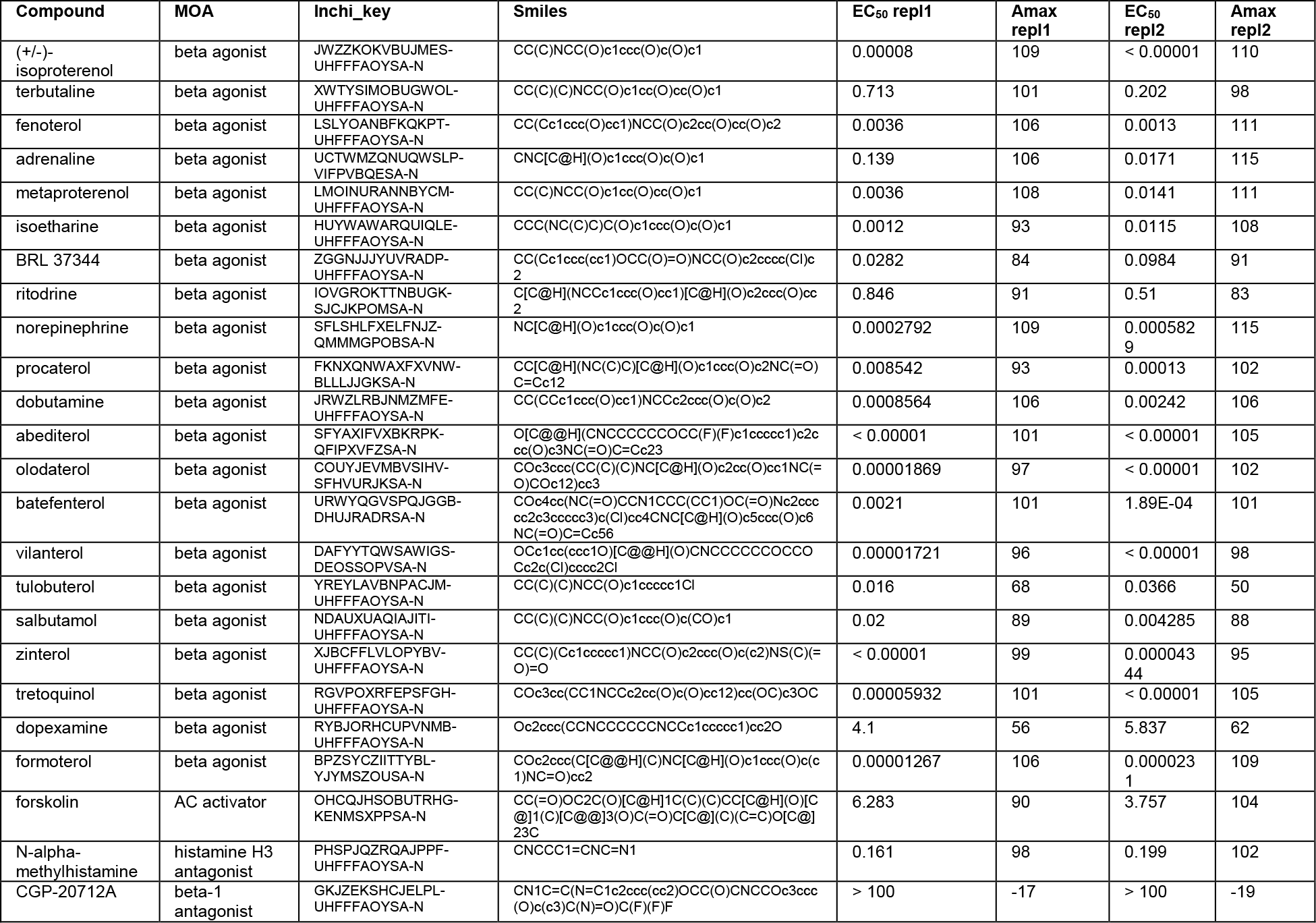
Overview over compounds used in THP1 experiments, comparing cAMP and QuantiGene readouts.

**Supplementary Figure 1:**
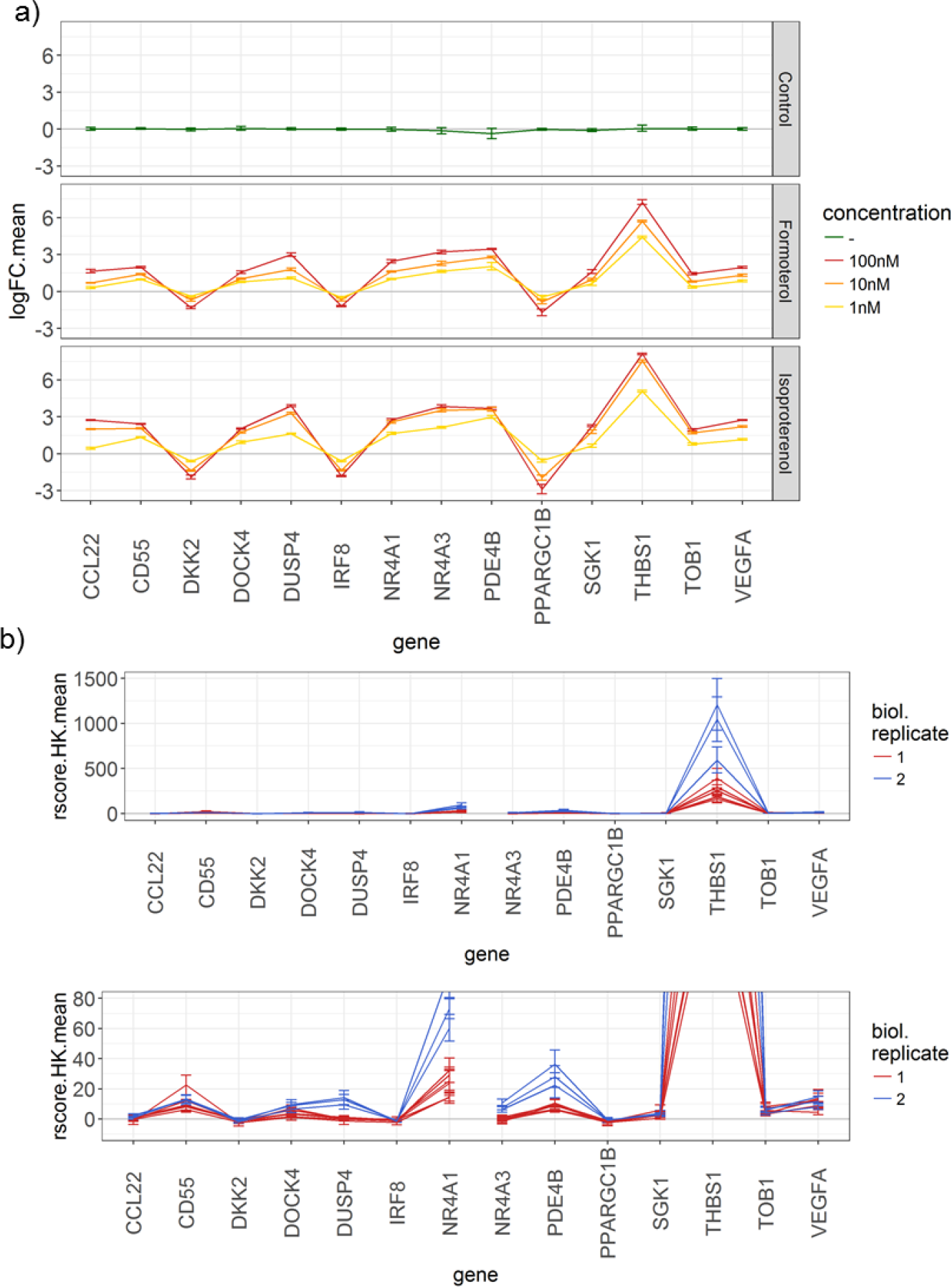
Validation of the beta agonists gene-signature. a) qPCR results for DMSO, formoterol and isoproterenol demonstrate dose dependent effect on genes after 4h incubation. b) QuantiGene Plex results for the two gene-signatures for 10uM of isoproterenol after 4h incubation, shown at two different scales: upper = full scale with THBS1 having a much stronger response than the other genes, and lower = y-axis cut at 80 to visualize the genes with lower variance. Shown are mean standard deviations of the rscore_HK values of active control wells of each plate.

**Supplementary Figure 2:**
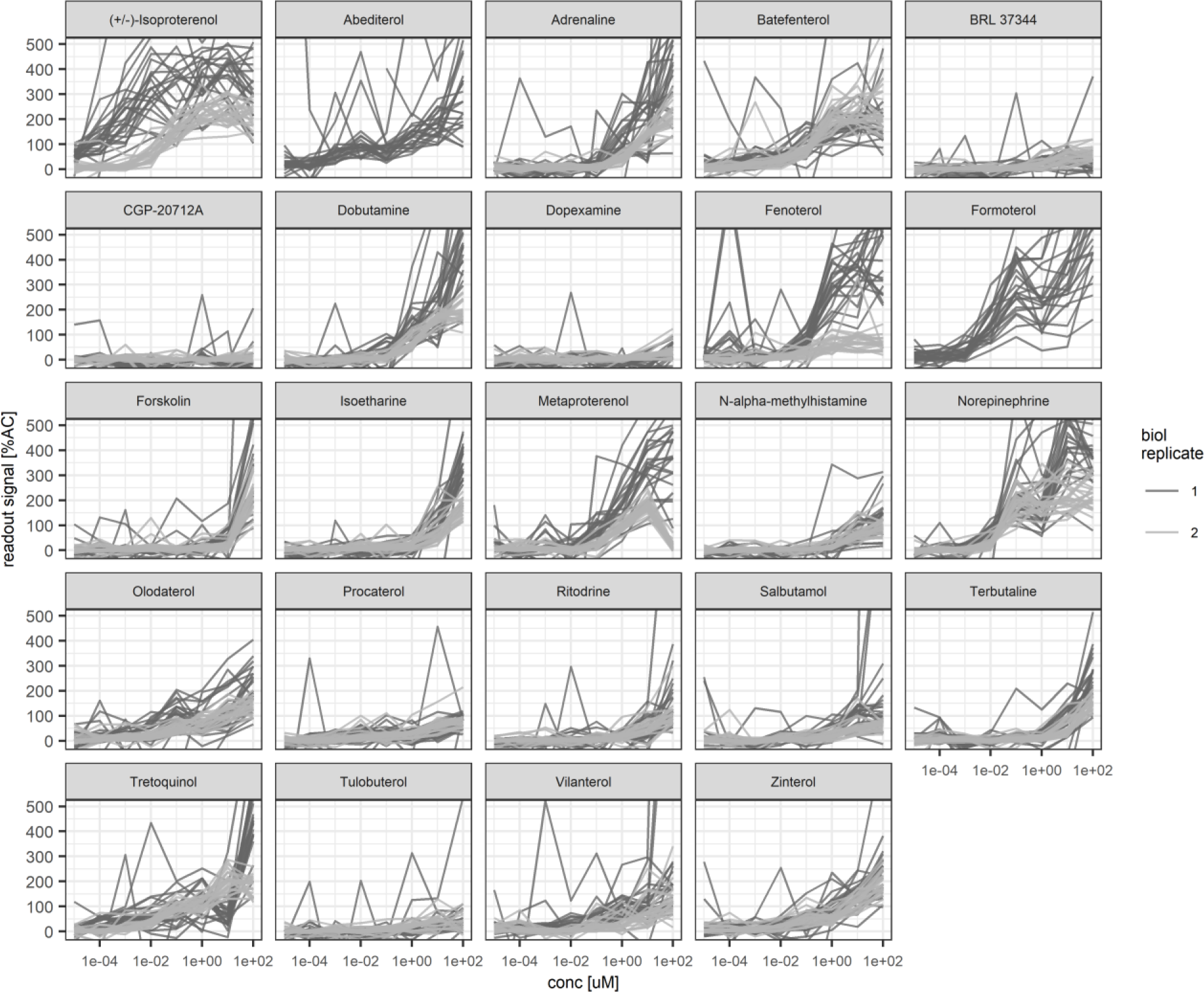
Dose-response of genes for each compound in the beta agonists dataset.

**Supplementary Figure 3:**
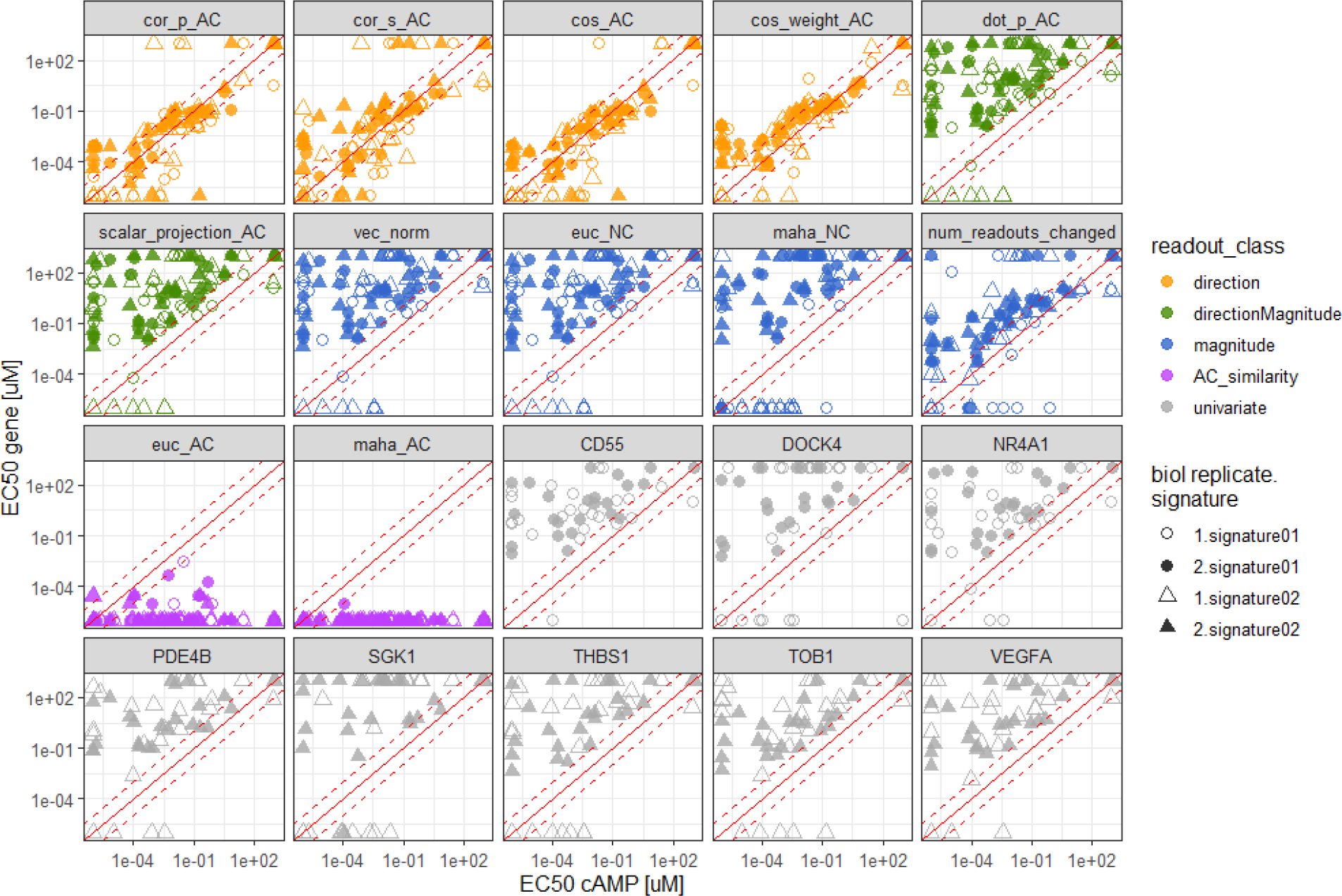
Correlation of gene and gene-signature EC_50_s with cAMP EC_50_s of the beta agonist dataset.

**Supplementary Figure 4:**
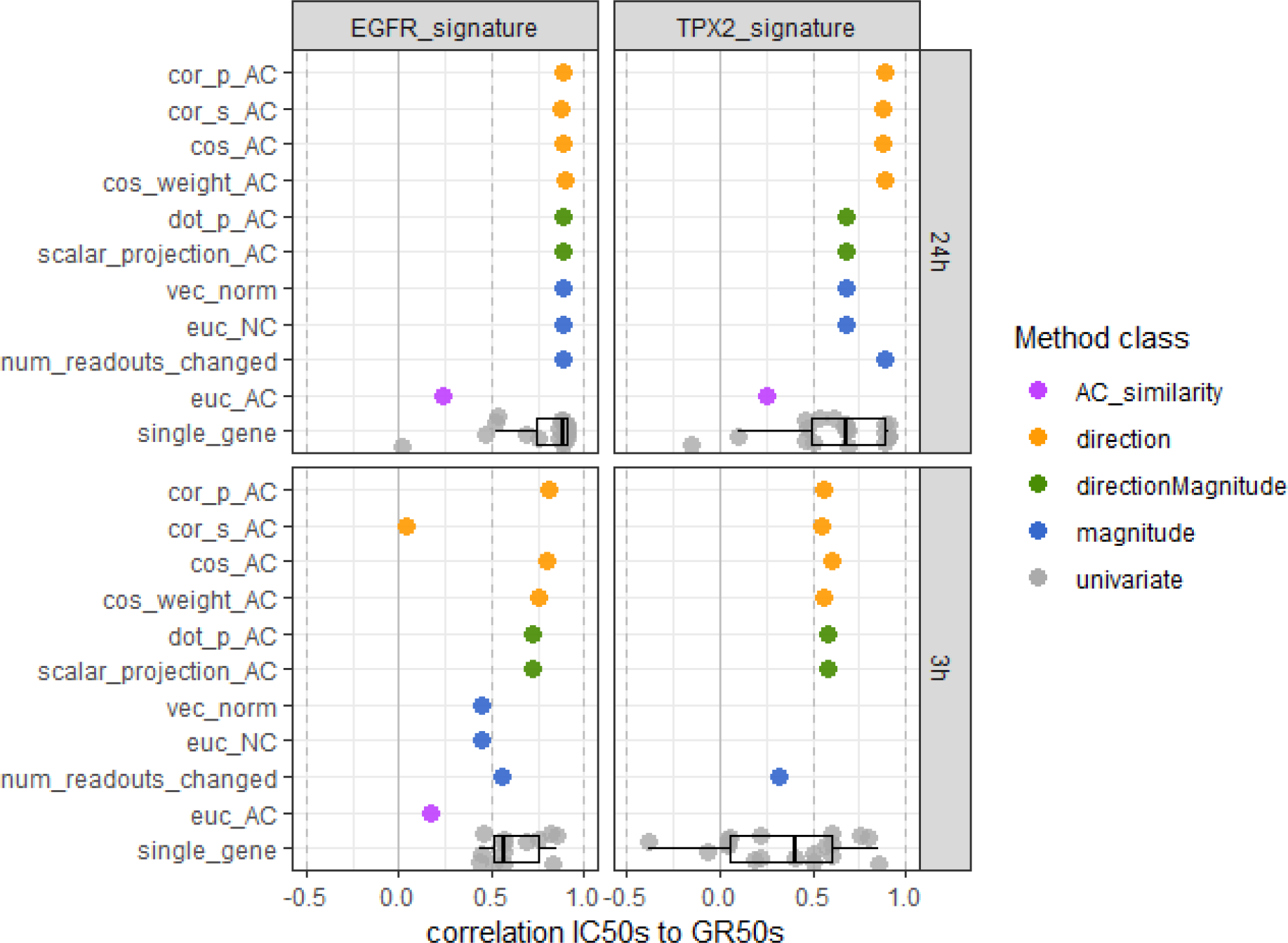
Correlation of cell growth inhibition GR_50_s with all gene and gene-signature EC_50_s of the EGFR inhibitor dataset.

**Supplementary Figure 5:**
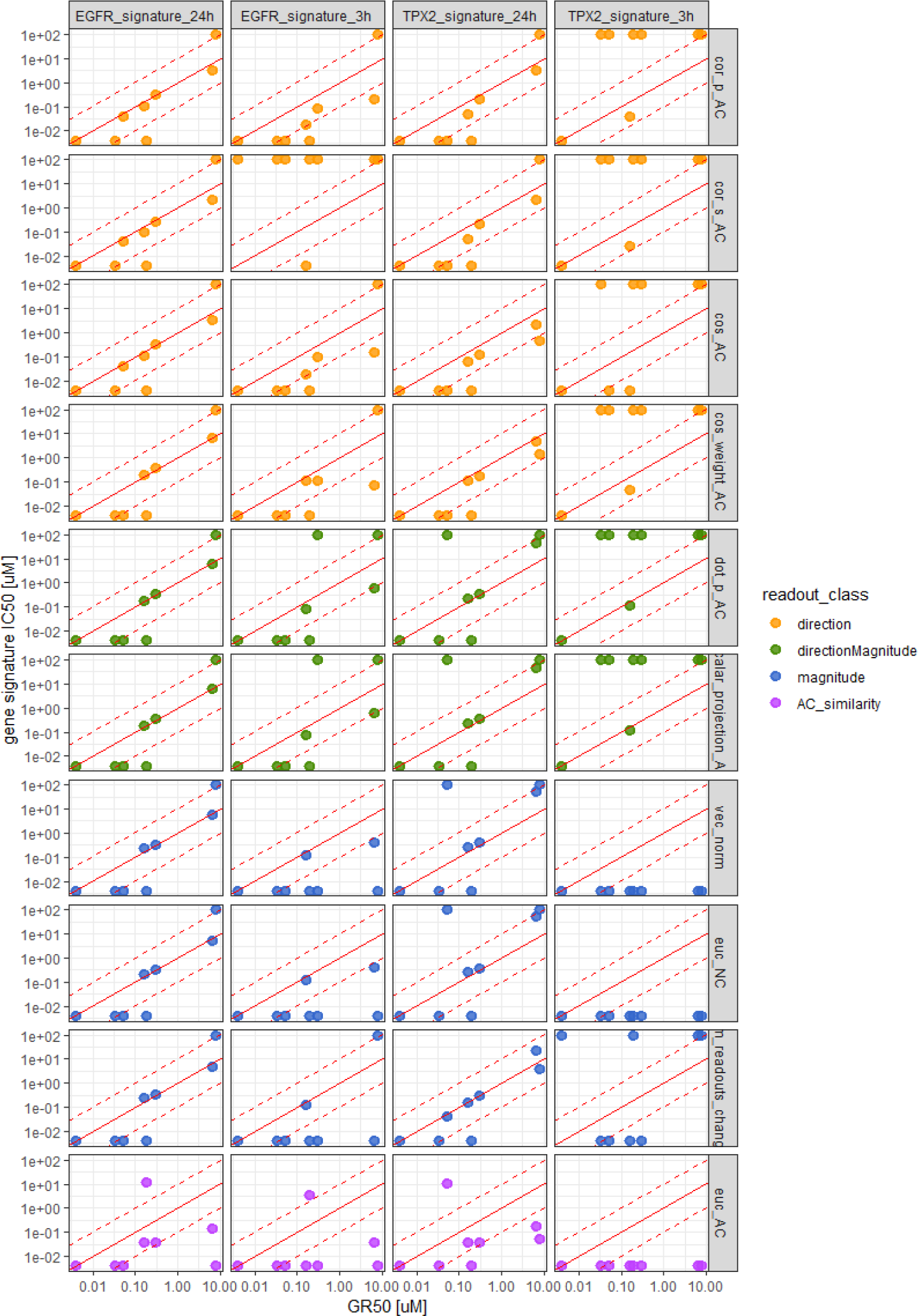
Correlation of gene-signature IC_50_s to GR_50_s of the EGFR inhibitor dataset.

**Supplementary Figure 6:**
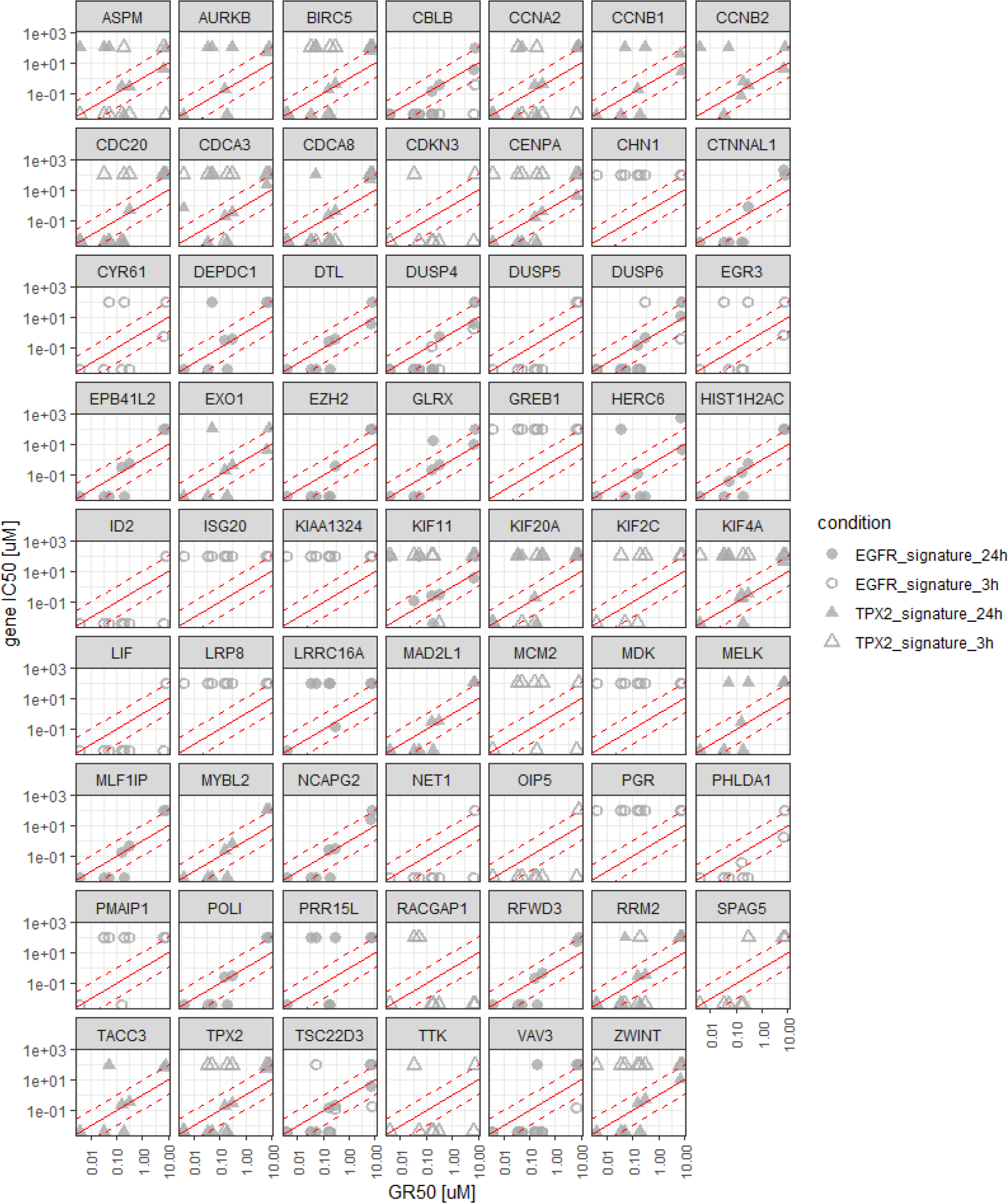
Correlation of single gene IC_50_s to GR_50_s of the EGFR inhibitor dataset, for the 20 genes most responding to the active control from the EGFR and TPX2 signatures.

## References

1. Perou, C.M., et al., Molecular portraits of human breast tumours. Nature, 2000. 406(6797): p. 747–52.

2. Sotiriou, C., et al., Gene expression profiling in breast cancer: understanding the molecular basis of histologic grade to improve prognosis. J Natl Cancer Inst, 2006. 98(4): p. 262–72.

3. Farmer, P., et al., A stroma-related gene signature predicts resistance to neoadjuvant chemotherapy in breast cancer. Nat Med, 2009. 15(1): p. 68–74.

4. Lamb, J., et al., The Connectivity Map: using gene-expression signatures to connect small molecules, genes, and disease. Science, 2006. 313(5795): p. 1929–35.

5. Scherf, U., et al., A gene expression database for the molecular pharmacology of cancer. Nat Genet, 2000. 24(3): p. 236–44.

6. Li, H., J. Qiu, and X.D. Fu, RASL-seq for massively parallel and quantitative analysis of gene expression. Curr Protoc Mol Biol, 2012. Chapter 4: p. Unit 4 13 1–9.

7. Ye, C., et al., DRUG-seq for miniaturized high-throughput transcriptome profiling in drug discovery. Nat Commun, 2018. 9(1): p. 4307.

8. Guibert, N., et al., Amplicon-based next-generation sequencing of plasma cell-free DNA for detection of driver and resistance mutations in advanced non-small cell lung cancer. Ann Oncol, 2018. 29(4): p. 1049–1055.

9. Xu, C., et al., Detecting very low allele fraction variants using targeted DNA sequencing and a novel molecular barcode-aware variant caller. BMC Genomics, 2017. 18(1): p. 5.

10. Bush, E.C., et al., PLATE-Seq for genome-wide regulatory network analysis of high-throughput screens. Nat Commun, 2017. 8(1): p. 105.

11. Subramanian, A., et al., A Next Generation Connectivity Map: L1000 Platform and the First 1,000,000 Profiles. Cell, 2017. 171(6): p. 1437–1452 e17.

12. Chen, M.H., et al., Gene expression-based chemical genomics identifies potential therapeutic drugs in hepatocellular carcinoma. PLoS One, 2011. 6(11): p. e27186.

13. De Wolf, H., et al., High-Throughput Gene Expression Profiles to Define Drug Similarity and Predict Compound Activity. Assay Drug Dev Technol, 2018. 16(3): p. 162–176.

14. Hahn, C.K., et al., Proteomic and genetic approaches identify Syk as an AML target. Cancer Cell, 2009. 16(4): p. 281–94.

15. Hahn, C.K., et al., Expression-based screening identifies the combination of histone deacetylase inhibitors and retinoids for neuroblastoma differentiation. Proc Natl Acad Sci U S A, 2008. 105(28): p. 9751–6.

16. Peck, D., et al., A method for high-throughput gene expression signature analysis. Genome Biol, 2006. 7(7): p. R61.

17. Stegmaier, K., et al., Gene expression-based high-throughput screening(GE-HTS) and application to leukemia differentiation. Nat Genet, 2004. 36(3): p. 257–63.

18. House, J.S., et al., A Pipeline for High-Throughput Concentration Response Modeling of Gene Expression for Toxicogenomics. Front Genet, 2017. 8: p. 168.

19. Hu, J., et al., Analysis of dose-response effects on gene expression data with comparison of two microarray platforms. Bioinformatics, 2005. 21(17): p. 3524–9.

20. Ji, R.R., et al., Transcriptional profiling of the dose response: a more powerful approach for characterizing drug activities. PLoS Comput Biol, 2009. 5(9): p. e1000512.

21. Lin, D., et al., Classification of Trends in Dose-Response Microarray Experiments Using Information Theory Selection Methods. The Open Applied Informatics Journal, 2009(3): p. 34–43.

22. Lin, D., et al., Testing for trends in dose-response microarray experiments: a comparison of several testing procedures, multiplicity and resampling-based inference. Stat Appl Genet Mol Biol, 2007. 6: p. Article26.

23. Pramana, S., et al., IsoGene: An R Package for Analyzing Dose-response Studies in Microarray Experiments. The R Journal, 2010. 2(1): p. 5–12.

24. Duan, Q., et al., L1000CDS(2): LINCS L1000 characteristic direction signatures search engine. NPJ Syst Biol Appl, 2016. 2.

25. Gabriel, D., et al., High throughput screening technologies for direct cyclic AMP measurement. Assay Drug Dev Technol, 2003. 1(2): p. 291–303.

26. Hafner, M., et al., Growth rate inhibition metrics correct for confounders in measuring sensitivity to cancer drugs. Nat Methods, 2016. 13(6): p. 521–7.

27. Farmer, P. and J. Pugin, beta-adrenergic agonists exert their “anti-inflammatory” effects in monocytic cells through the IkappaB/NF-kappaB pathway. Am J Physiol Lung Cell Mol Physiol, 2000. 279(4): p. L675–82.

28. Creighton, C.J., et al., Activation of mitogen-activated protein kinase in estrogen receptor alpha-positive breast cancer cells in vitro induces an in vivo molecular phenotype of estrogen receptor alpha-negative human breast tumors. Cancer Res, 2006. 66(7): p. 3903–11.

29. Abraham, Y., X. Zhang, and C.N. Parker, Multiparametric Analysis of Screening Data: Growing Beyond the Single Dimension to Infinity and Beyond. J Biomol Screen, 2014. 19(5): p. 628–39.

30. Loo, L.H., L.F. Wu, and S.J. Altschuler, Image-based smultivariate profiling of drug responses from single cells. Nat Methods, 2007. 4(5): p. 445–53.

31. Liberzon, A., et al., Molecular signatures database (MSigDB) 3.0. Bioinformatics, 2011. 27(12): p. 1739–40.

32. Subramanian, A., et al., Gene set enrichment analysis: a knowledge-based approach for interpreting genome-wide expression profiles. Proc Natl Acad Sci U S A, 2005. 102(43): p. 15545–50.

33. Gubler, H., et al., Helios: History and Anatomy of a Successful In-House Enterprise High-Throughput Screening and Profiling Data Analysis System. SLAS Discov, 2018. 23(5): p. 474–488.

